# Effects of selection for early dispersal on the ambrosia beetle *Xyleborinus saxesenii* and its fungal symbionts

**DOI:** 10.1101/2024.07.30.605773

**Authors:** Antoine Melet, Peter Biedermann

## Abstract

Overlapping generations is a defining characteristic of advanced social life. In cooperative breeding societies, for example, temporary groups of mature offspring are formed that assist in the rearing of additional brood before the offspring disperse and reproduce independently. It is hypothesized that the number of helpers and their delayed dispersal period will determine the number of siblings that can be reared, thus resulting in an indirect fitness gain. The objective of this study, was to investigate the effect of artificial selection for early dispersal of mature offspring on the life history, behaviour and fungal symbionts in the cooperatively breeding ambrosia beetle *Xyleborinus saxesenii*. Two lineages of beetles were bred in the laboratory for five successive generations. In one group, dispersing females were selected at random to initiate the next generation, while in the other group, only early dispersers were selected. A number of life-history traits exhibited a pronounced response in the initial generation, subsequently recuperating to levels approximating those observed at the outset of the experiment. The laboratory rearing resulted in an increasing proportion of successful nests in both lineages. Additionally, the control lineage exhibited a reduction in lifespan and in productivity. Furthermore, significant differences were observed in the fungal communities from the third generation onwards. The results suggest that *X. saxesenii* has limited potential to respond to this selection pressure, potentially due to sibmating and resulting low genetic variability. Furthermore, the correlation between nest lifespan and productivity is a crucial factor in explaining philopatry and altruism in this species.

## Introduction

A multitude of animal species live in groups, ranging from the relatively simple herds of bison to the highly advanced hives of honeybees. Advanced sociality is defined by a number of characteristics, including a high reproductive skew, the frequent occurrence of alloparental care and a high level of relatedness between helpers, reproducers and offspring (Wilson, 1971). The most derived social lifestyles are cooperative breeding and eusociality. In these societies, offspring typically do not disperse immediately after maturation. Instead, they overtake cooperative tasks in the natal nest. either for a limited period (cooperative breeding) or permanently (eusociality). It is therefore not surprising that alloparental care has evolved alongside philopatry in such societies (Hochberg et al., 2008; Le Galliard et al., 2005; Mullon et al., 2018). This theoretical assumption is also supported by correlative evidence in a wide range of social organisms ranging from microbes to vertebrates (Choe & Crespi, 1997; Fisher et al., 2013; Hamilton, 1964; Hatchwell, 2009; Koenig & Dickinson, 2016; Rainey & Rainey, 2003; Taborsky, 1994). Nevertheless, there is currently a paucity of experimental evidence to support the co-evolution of philopatry and alloparental care.

*Xyleborinus saxesenii* Razteburg (Curculionidae: Scolytinae: Xyleborini) is a Eurasian ambrosia beetle that lives in cooperatively breeding societies with variable durations of female offspring philopatry and alloparental care (Biedermann & Taborsky, 2011). Adult female offspring typically delay dispersal following fertilization by their brothers. While residing in their natal nest, daughters assist their mother with brood care, fungus farming and the enlargement of tunnels (Biedermann & Taborsky, 2011; Peer & Taborsky, 2007). In *Xyleborus affinis* Eichhoff, a related species with a similar social lifestyle, a long philopatric time has been demonstrated to be associated with a direct fitness cost later in life (Biedermann et al., 2011). Both species can be reared in an artificial medium (Biedermann et al., 2009), which provides the unique possibility of selecting short philopatric periods of adult offspring and consequently testing for correlated effects on alloparental care.

Ambrosia beetles are typically associated with mutualistic fungi, which they cooperatively cultivate in monoculture within tunnel systems and which serve as food for both larvae and adults (Biedermann & Vega 2020). *Xyleborinus saxesenii* cultivates two nutritional fungi that develop in succession, *Dryadomyces sulphureus* and *Raffaelea canadensis* (Biedermann et al., 2013; Diehl et al., 2022; Francke-Grosmann, 1975). Other fungal species are also present within the nests of *X. saxesenii,* although they are not as closely associated with the beetle (Biedermann et al., 2013). However, the accumulation of less beneficial fungi within nests over time also affects the fitness of late-dispersing offspring, which vertically transmit all these fungi to newly founded nests (Diehl et al., 2022). A strong correlation has been identified between highly productive nests, their durability, long philopatric periods and alloparental care by adult daughters (Biedermann & Taborsky 2011, Nuotclá et al. 2021). It has therefore been hypothesized that the management of beneficial fungal communities is a significant factor in the durability of nests and, consequently, in the philopatry in ambrosia beetles (Biedermann & Rohlfs 2017).

Artificial selection experiments represent a valuable research tool for the study of evolutionary processes. Phenotypic responses can be measured as a direct response to selective pressure (Brakefield, 2003; Conner, 2003). In natural conditions, adaptation is typically too slow to be observed within the span of a scientific project or even a lifetime. By imposing strong selection pressures in controlled conditions, researchers can detect adaptive changes over a period of just a few generations (Conner, 2003). When combined with the relatively short generation time of certain insects, it is possible to observe adaptation over a period of just a few months. The use of artificial selection experiments allows for a direct experimental test of hypotheses, which can be used to complement the results of correlative studies (Lewis & Morran, 2022). Such experiments are also used to study communities of two or multiple species (Blouin et al., 2015; Swenson et al., 2000).

The objective of this study was to examine the relationship between brief philopatric periods in adult female offspring and the expression of other life history traits in the subsequent generation of offspring, with a particular focus on alloparental care and nest founding success. It was hypothesized that beetles selected for early dispersal would exhibit a tendency to disperse earlier and earlier over generations, produce less-durable nests with fewer offspring and demonstrate a reduction in alloparental care behaviors. Furthermore, we sought to ascertain the impact of offspring philopatry has on fungal communities. It was hypothesized that these communities would be less beneficial in nests of females with long philopatric periods and that they would contain higher relative abundances of *R. canadensis* than *D. sulphureus* fungal cultivars (Diehl et al., 2022).

## Material and methods

### Model species

*Xyleborinus saxesenii* females emerge from their native nest already mated and proceed to initiate new nests by excavating in suitable pieces of wood. They tunnel deep into the xylem, inoculating the wood with mutualistic fungi that they have transported from their native nest in mycetangia (specialized organs for transporting fungal spores) and guts (Batra, 1966; Francke-Grosmann, 1975; Mayers et al., 2022). Once the fungi are established, the foundress begins to lay eggs. The mutualistic fungi serve as the food source for the adults and to a major extent also the larvae (i.e. larvae feed on fungus-infested xylem; De Fine Licht & Biedermann, 2012).

### Artificial rearing

*Xyleborinus saxesenii* was bred in transparent plastic tubes filled to two-third capacity with artificial rearing media based on beech sawdust, agar and additional nutrients (see standard media in Biedermann et al. 2009). All nests in this experiment were kept at a constant temperature of 25°C, 70% humidity and constant darkness. Thirty nests from the breeding system were randomly selected to serve as the basis for this experiment (F0). The nests were randomly assigned to the control or the treatment lineage (control and early dispersers, respectively). The dispersing females from F0 were collected, briefly washed in 70% ethanol and then rinsed in distilled water to remove the ethanol. The primary fungal mutualists are not adversely affected by this protocol of sterilization (Biedermann et al. 2009) since they are protected within the mycetangia. Subsequently, the dispersing females were briefly dried on sterile tissue paper and immediately singly transferred to rearing medium within plastic tubes that were closed with plugs of foamed plastic. All nests were individually labelled and their founding date recorded. The nests were observed three times a week until they ended.

Pilot studies have indicated that periods of diapause are essential for the successful long-term breeding of *X. saxesenii*. Consequently, breeding temperature was reduced during the second generation (F2) for a ten-weeks period to 8°C, in order to simulate the overwintering period. Egg-laying ceased during this period. Only larvae and adults were observed in the diapausing galleries, along with dead pupae that were cannibalized rapidly at the end of the diapause period. No dispersal was observed during the diapause period.

### Directional artificial selection

For each generation of the treatment lineage, 30 nests were randomly selected. From each nest, the seven initial dispersing females were collected, washed, rinsed and dried as previously described. They were then placed individually in tubes containing fresh medium to initiate the subsequent generation. In each generation of the control lineage, 30 nests were randomly selected. The dispersers were collected from these nests in order to initiate the subsequent generation. The specimens were washed, rinsed and dried, then placed in tubes as previously described. Once all dispersers from a nest had been collected, the first 25% and last 25% of nests descending from that particular parent nest were removed. This was essential to prevent temporal overlap in the dispersal periods. The selection regime was maintained over five successive generations (i.e. treatment F1 to F5 and control F1 to F5).

The experimental design yielded ten experimental groups. The objective was to study the effect of the selection at every step, by comparing consecutive generations within each lineage (control F1 vs. F2, F2 vs. F3, F3 vs. F4, F4 vs. F5 and treatment F1 vs. F2, F2 vs. F3, F3 vs. F4, F4 vs. F5), the ultimate effect of the selection, by comparing the first and last generation within each lineage (control F1 vs. F5 and treatment F1 vs. F5), and the divergence between the two lineages, by comparing the two groups within each generation (control F1 vs. treatment F1, control F2 vs. treatment F2, control F3 vs. treatment F3, control F4 vs. treatment F4, control F5 vs. treatment F5). Particular attention was paid to the initial generation, with the objective of identifying changes that occur at an early stage of the selection process.

### Success rate

To evaluate the success rate of each group, we counted the nests that produced at least two dispersing females as successful ones. This approach was employed because, with nests that produced only one dispersing female, it is not possible to be sure that the single female was not the foundress, leaving her nest. The success rates were compared using pairwise Chi-square tests, with the Holm correction applied to account for multiple testing.

### Nest development

The number of eggs, larvae, pupae, adult female in the nest and dispersing females was recorded for each observation. The three larval instars were not distinguished. A nest was deemed to have reached the end of its lifespan if no adults and no dispersers were observed for a period of seven consecutive days. The final observations, which did not include any record of adults or dispersers, were excluded from the dataset to prevent the lifespan of nests from being artificially inflated. From the census data, we extracted two key variables: (i) the total lifespan of the nest, defined as the number of days between the starting day and the day the last disperser was observed, and (ii) the timing of first dispersal, defined as the number of days between the starting day and the day the first disperser was observed.

The developmental times were compared using pairwise Wilcoxon tests, with the Holm correction to account for multiple testing. The dispersal patterns were compared with pairwise survival analysis, which was based on a Cox proportional hazards model, and followed by a multiple comparison of means using Tukey contrasts.

### Productivity

The total number of dispersers collected from a nest was used as the productivity of said nest. The productivities were compared using pairwise Wilcoxon tests, with the Holm correction to account for multiple testing. The effect of the timing of first dispersal on the productivity was analysed using a linear model.

### Behavioral observations

Following each census count of a nest, a scan observation was conducted, whereby the behavior of each encountered larva and adult was recorded in accordance with the method outlined by Biedermann & Taborsky (2011). The behaviors exhibited by larvae and adults were recorded separately. The duration of scan observations was kept to a minimum in order to obtain a snapshot overview the behavior exhibited by the entire nest. No behavioral data were recorded between the date of 11.07.2022 and 18.07.2022, and between 08.08.2022 and 15.08.2022. From the behavioral records, two categories of behaviors were analyzed: (i) activity, defined as all the behaviors except resting behavior, and (ii) social behaviors, defined as the behaviors directed towards nest excavation and hygiene (i.e. digging, balling, grooming and cannibalism for the larvae and digging, shuffling, grooming and cannibalism for the adults).

The activity and social behaviors of larvae and adult females were analysed separately with multi-level generalized linear models specifying binomial errors, with the nest identity fitted as a random effect, followed by a multiple comparison using Tukey contrasts.

### Fungal community sampling and DNA extraction

From the second generation onwards, nests were randomly selected from each group. The seven initial dispersing females from the selected nests were collected and frozen at –20° C until DNA extraction and fungal community sequencing. Some nests did not yield any dispersers, resulting in the following final sample sizes for the different groups: control F2 (N = 8), F3 (N = 8), F4 (N = 8), F5 (N = 8), and treatment F2 (N = 7), F3 (N = 6), F4 (N = 7), F5 (N = 7). The first generation was not included due to a lack of available nests. Nests selected for the fungal community analysis were excluded from the life history and behavioral analyses.

In order to extract fungal DNA, the seven females that dispersed from a nest were combined into a single sample. The samples were subjected to mechanical grinding at 2700 rpm for 20 minutes in a ZR BashingBead Lysis Tube 2,0 mm with 750 µL of lysis solution (Zymo Research, Germany) on a Vortex Genie 2 (Scientific Industries). After centrifugation at 18 000 g for 1 min, the supernatant was collected and transferred to a ZR BashingBead Lysis Tube 0.1 and 0.5 mm. Thereafter, 300 µL of lysis solution were added, and the mixture was ground at 2700 rpm for 20 minutes on a Vortex Genie 2. Subsequently, DNA was extracted utilising ZymoBIOMICS DNA Miniprep kits (Zymo Research, Germany) in accordance with the manufacturer’s instructions. The isolated DNA was stored at –20° C until processing.

### Amplicon sequencing of fungal communities

The amplification of fungal DNA was achieve through the use of LSU (28S) rRNA primers, as the primers typically employed for the analysis of fungal communities, targeting the ITS region, proved ineffective in amplifying Ophiostomataceae species, including *Raffaelea* and *Dryadomyces* (Kostovcik et al., 2015). The PCR conditions were as follows: an initial denaturation at 98°C for 3 minutes, 35 cycles of denaturation at 98°C for 10 seconds, annealing at 54°C for 30 seconds and elongation at 72°C for 20 seconds; followed by a final extension step at 72°C for 10 minutes.

Following amplification, the DNA samples were sequenced by the company StarSeq (Mainz, Germany) using the primers described, on the Illumina MiSeq v3 2 x 300-bp platform in accordance with the Illumina protocol. To ensure the quality of our fungal community results, we included negative and positive controls alongside our samples. The negative controls were composed of DNA extracts that have been prepared and sterilized as previously described, yet lacked the presence of the beetles. Two negative controls were prepared, alongside the samples from the groups control F5 and treatment F5 respectively. Furthermore, the sequencing company utilised two samples of pure water, which were devoid of any DNA, and which underwent all library preparation steps. The positive control was a mock community that had been prepared for another experiment and sent for sequencing within the same batch. The mock community comprised equal amounts of *Dryadomyces sulphureus, Raffaelea canadensis, Beauveria bassiana* and the yeasts *Pichia sp.* and *Candida sp.* DNA was extracted as previously described. The positive control provided by the sequencing company comprised a defined quantity of DNA from eight bacterial species (*Pseudomonas aeruginosa, Escherichia coli, Salmonella enterica, Lactobacillus fermentum, Enterococcus faecalis, Staphylococcus aureus, Listeria monocytogenes, Bacillus subtilis*) and two yeast species (*Saccharomyces cerevisiae, Cryptococcus neoformans*) (ZymoBIOMICS Microbial community DNA standard, Zymo Research, Germany). The mock community and the positive control provided by the company were replicated twice, resulting in a total of four positive controls.

### Bioinformatics processing

The raw, demultiplexed reads were processed using Usearch v11 (Edgar, 2010). The forward and reverse reads were merged using the *-fastq_mergepairs* command, with a minimum of 200 base pairs for the merged sequence and a maximum of 20 mismatches in the alignment as a preliminary quality filtering step. The primers were removed from the reads using the *-fastx_truncate* command, and the overall quality was then assessed using the *-fastq_filter* command, with a total expected error threshold of 1. Unique sequences were identified using the *-fastx_uniques* command and the singletons were excluded using the –*sortbysize* command with a minimum size of 2. The amplicon reads were denoised using the –*unoise3* command. This command does not cluster similar sequences; rather, it identifies and corrects reads with sequencing errors and removes chimeras, resulting in amplicon sequence variants (ASVs), a higher-resolution analogue of the traditional OTU (Edgar, 2016b). The ASVs were taxonomically classified in two stages using the *-usearch_global* command. The command is based on the USEARCH algorithm, which searches for high-identity hits to a database sequence (Edgar, 2010). In the initial stage, a LSU sequences database of stock culture of fungi isolated from *X. saxesenii* was employed. The database comprised eighteen reference sequences of twelve fungal species (Diehl et al., 2022). The identity threshold was set at 97% due to the fact that the database comprises sequences of known fungal symbionts of *X. saxesenii.* The unclassified ASVs were employed as the input for the second stage, which utilised a custom reference database constructed from NCBI data using BCdatabaser (Keller et al., 2020). The second database comprised 82 250 sequences (Diehl et al., 2022). In order to ensure accurate classification, the identity threshold was set at 99%. In instances where the taxonomic outputs differed between the two stages, the output from the initial stage was retained. The remaining unclassified ASVs were then used as input for the final stage, emplying the *-sintax* command with a sintax cutoff of 0.8. The SINTAX algorithm is comparable to a naïve Bayesian classifier algorithm but it does not require training (Edgar, 2016a).

### Statistical analysis of metabarcode data

In order to enhance the quality of our dataset, we employed a contaminant removal method in accordance with the R package ‘decontam’, taking the negative controls into account. This process reduces the complexity of the microbiome data in downstream analysis while preserving their integrity (Davis et al., 2018). The positive and negative controls were visualized but excluded from the sample set. Following decontamination and the exclusion of controls, the samples were rarefied, accounting for unequal numbers of reads between samples. The sample with the fewest reads had 9 838 reads; this threshold was used to generate the rarefied samples, which were subjected to subsequent analysis.

The alpha diversity of the fungal communities was compared between the various groups by means of a comparison of the observed ASVs richness and Shannon’s diversity index. Pairwise Wilcoxon tests were employed to compare the groups, with Bonferroni correction to account for multiple testing. The beta diversity was analysed through the calculation of dissimilarity matrices using the Bray-Curtis method, considering both the presence/absence and relative abundances of ASVs. A pairwise PERMANOVA was used to analyze the matrices. Composition barplots were constructed for visualization, aggregated to the genus level and faceted by group.

### Statistical analysis

All tests were conducted using R version 4.0.2 (R Core Team 2020), with the RStudio interface version 1.3.1073. The additional packages *car* (Fox & Weisberg, 2019), *decontam* (Davis et al., 2018)*, lme4* (Bates et al., 2015), *multcomp* (Hothorn et al., 2008), *pairwiseAdonis* (Martinez Arbizu, 2020), *phyloseq* (McMurdie & Holmes, 2013), *rstatix* (Kassambara, 2023), *survival* (Therneau, Terry M. & Grambsch, Patrica M., 2000) and *vegan* (Oksanen et al., 2022) were employed.

## Results

Despite the objective of directional artificial selection being to accelerate female dispersal over five successive generations, no significant difference was observed in the time of dispersal between the two lineages (control F5 vs. treatment F5, Wilcoxon test, W = 3146, p = 0.18). Thus, the artificial selection pressure did not yield in the anticipated outcome. However, the data were duly reported and analyzed, in order to ascertain the reasons behind the failure of the artificial selection to produce the expected results.

### Success rate

In the control lineage, the success rate was 27% in F1, which was lower than in F5 (control F1 vs. F5; chi-square test, chi-squared = 28.494, p < 0.001) and in F2 (control F1 vs. F2; chi-square test, chi-squared = 41.472, p < 0.001). No other two successive generations were found to be significantly different (chi-squared tests, all p > 0.05). In the treatment lineage, the success rate was 24% in F1, which was lower than in F5 (treatment F1 vs. F5, chi-square test, chi-squared = 27.703, p < 0.001), but there was no significant difference in any of the successive generations (chi-squared tests, all p > 0.05). A comparison of the two lineages in each generation revealed no significant difference (chi-square tests, all p > 0.05) (Fig. 1).

**Figure 1:**
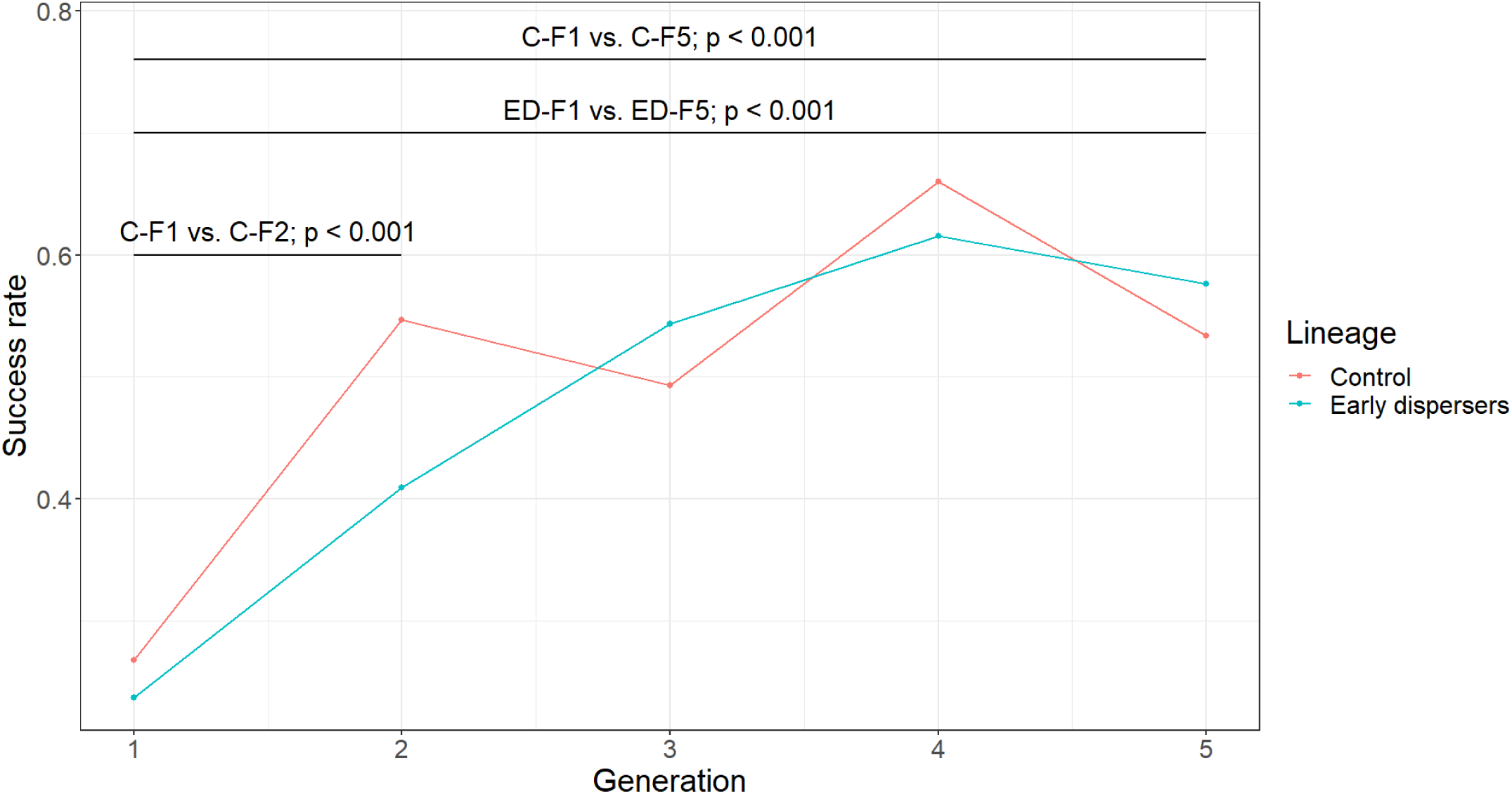
Success rate of the two lineages, from F1 to F5. The success rate is expressed as the proportion of nests that produced at least two dispersing females, out of the number of nests started. The comparisons were calculated using Chi-square tests, with the Holm correction for multiple testing applied to account for multiple testing

### Overall lifespan of nests

In the control lineage, the overall lifespan of nests was 51.13 days in F1, which was longer than in F5 (control F1 vs. F5, Wilcoxon test, W = 3876, p < 0.001), but there was no significant difference in any of the successive generations (Wilcoxon tests, all p > 0.05). In the treatment lineage, no two successive generations were found to be significantly different (Wilcoxon test, all p > 0.05). A comparison of the two lineages revealed a shorter lifespan in control F4 than in treatment F4 (control F4, vs. treatment F4, Wilcoxon test, W = 4174, p = 0.012), but the two lineages did not significantly differ in any of the other generations (Wilcoxon tests, all p > 0.05) (Fig. 2).

**Figure 2:**
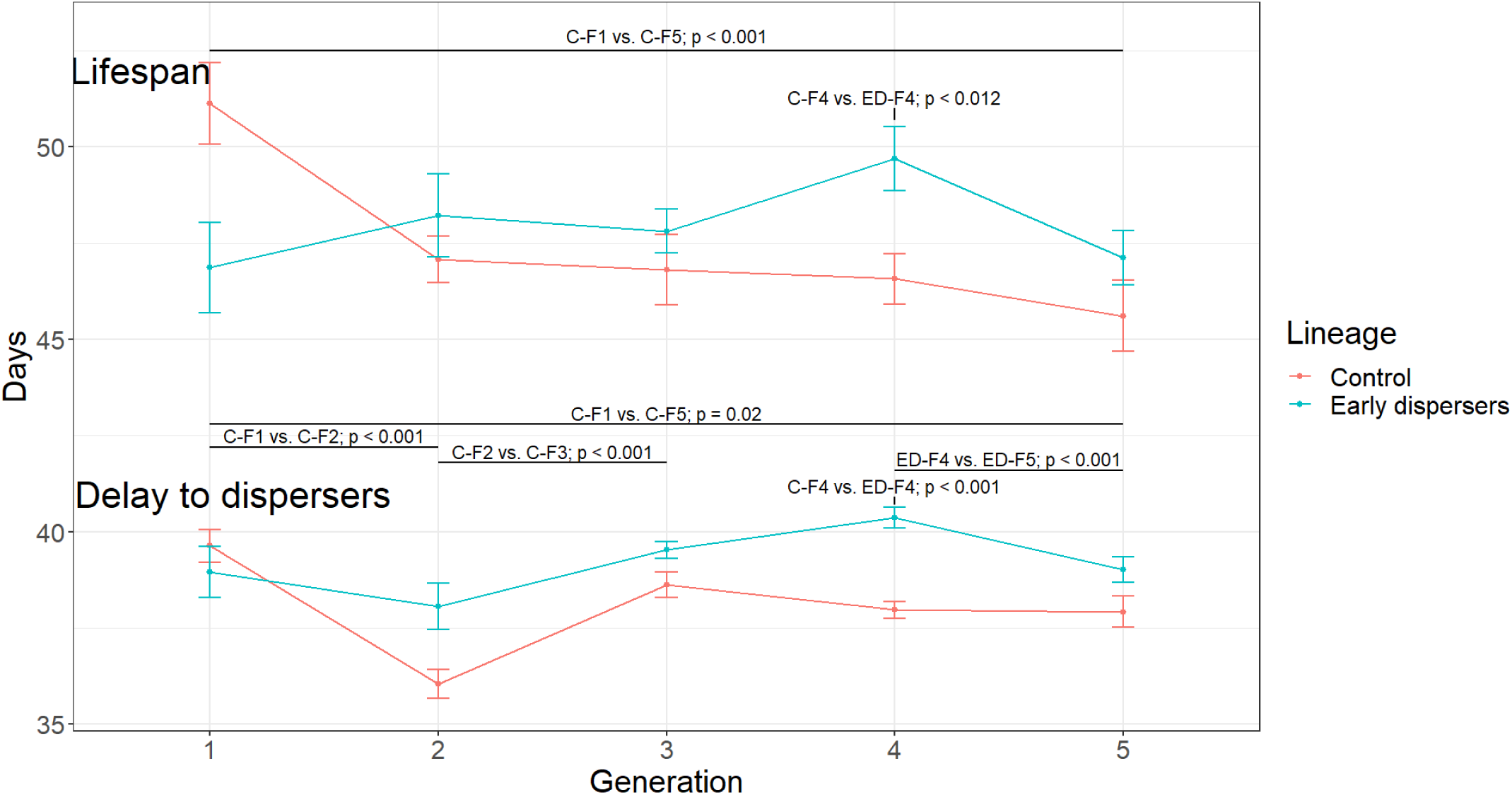
Development of the of the two lineages, from F1 to F5. The lifespan is expressed as the average number of days between the foundation of the nest and the dispersal of the last offspring. The delay to dispersers is the average number of days between the foundation of the nest and the dispersal of the first offspring. The comparisons were calculated with Wilcoxon tests including Holm correction for multiple testing. The vertical bars show the standard errors.

### Timing of first dispersal

In the control lineage, the duration before the start of dispersal was 39.63 days in F1, which was longer than in F5 (control F1 vs. F5, Wilcoxon test, W = 3596, p = 0.02). In the control lineage, the duration before the start of dispersal was longer in F1 than in F2 (control F1 vs. F2, Wilcoxon test, W = 7536, p < 0.001), and shorter in F2 than in F3 (control F2 vs. F3, Wilcoxon test, W = 3288, p < 0.001). In the control lineage, no other successive generations were found to be significantly different (Wilcoxon tests, all p > 0.05). In the treatment lineage, the duration before the start of dispersal was longer only in F4 than in F5 (treatment F4 vs. F5, Wilcoxon test, W = 6220, p < 0.001), and no other two successive generations were found to be significantly different (Wilcoxon tests, all p > 0.05). A comparison of the two lineages revealed only an earlier dispersal in treatment F4 than in control F4 (control F4 vs. treatment F4, Wilcoxon test, W = 2500, p < 0.001), but no differences in the other generations (Wilcoxon tests, all p > 0.05) (Fig. 2).

### Timing of overall dispersal

In the control lineage, overall dispersal occurred later in F1, than in F5 (control F1 vs. F5, Cox model, z = 4.264, p < 0.001) and in F2 (control F1 vs. F2, Cox model, z = 3.558, p = 0.012); no other two successive generations were found to be significantly different (Cox model, all p-values > 0.05). In the treatment lineage, no two successive generations were found to be significantly different (Cox models, all p > 0.05). A comparison of the two lineages revealed that overall dispersal occurred later in control F1 than in treatment F1 (control F1 vs. treatment F1, Cox model, z = 3.577, p = 0.012). The two lineages did not significantly differ in any of the other generations (Cox models, all p > 0.05) (Fig. 3).

**Figure 3:**
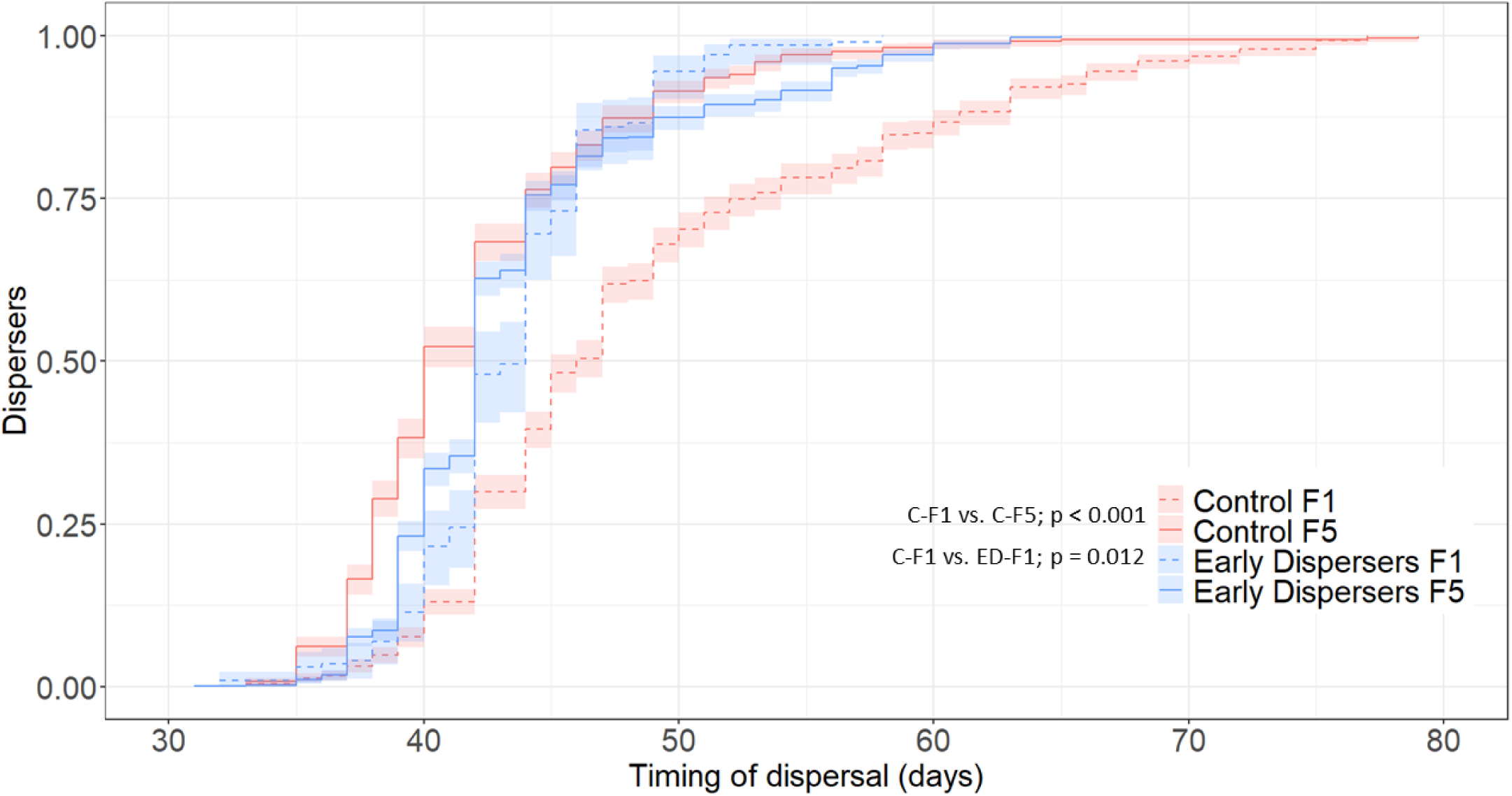
Dispersal pattern of the two lineages in generations F1 and F5. The dispersal pattern is expressed as a Cox proportional hazard model, with the dispersal of each offspring serving as the event. The timing of dispersal is expressed as the number of days between the foundation of the nest and the dispersal of an offspring. Colored areas indicate the confidence intervals.

### Productivity

In the control lineage, the productivity in F1 was not different from F5 (control F1 vs. F5, Wilcoxon test, W = 3308, p = 0.543), but higher than in F2 (control F1 vs. F2, Wilcoxon test, W = 7107, p < 0.001). Also, the productivity was lower in F2 than in F3 (control F2 vs. F3, Wilcoxon test, W = 3389, p = 0.002). In the control lineage, no other two successive generations were found to be significantly different (Wilcoxon tests, all p > 0.05). In the treatment lineage, the productivity was lower only in F2 than in F3 (treatment F2 vs. F3, Wilcoxon test, W = 1096, p < 0.001), but no other two successive generations were found to be significantly different (Wilcoxon test, all p > 0.05). A comparison of the two lineages revealed no significant difference in any of the generations (Wilcoxon tests, all p > 0.05) (Fig. 4).

**Figure 4:**
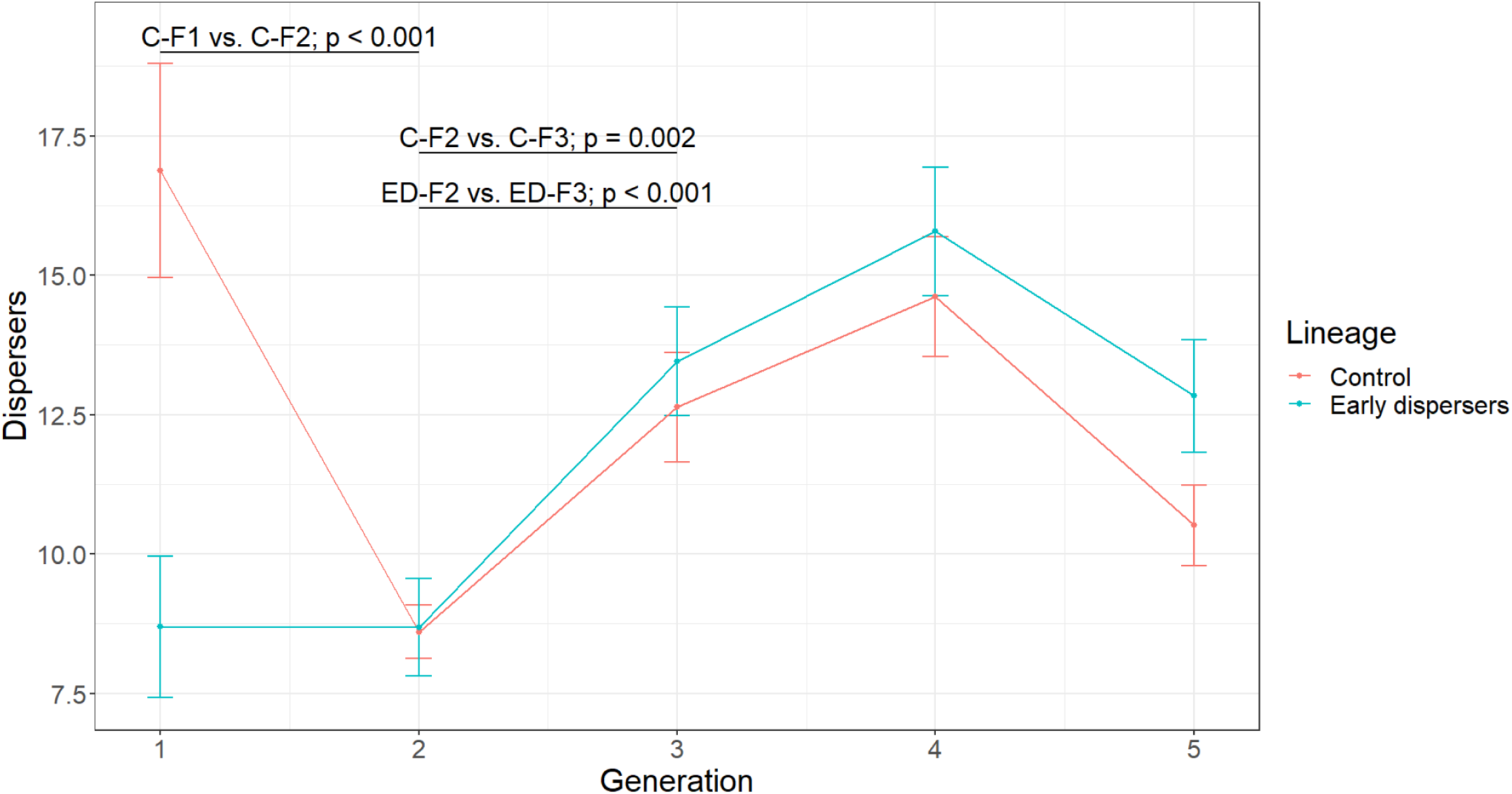
Productivity of the of the two lineages, from F1 to F5. The productivity is expressed as the average number of females that dispersed from the nests. The comparisons were calculated with Wilcoxon tests including Holm correction for multiple testing. The vertical bars show the standard errors.

Over all groups, the productivity was strongly correlated with the timing of dispersal (linear model, z = 5.55, p < 0.001).

### Behavioral observations

In both the control and the treatment lineage, the proportion of social behaviors of larvae was equal over all consecutive generations (linear models, all p > 0.05). There was no difference between the two lineages in any of the generations (linear models, all p > 0.05) (Fig. 5A).

**Figure 5:**
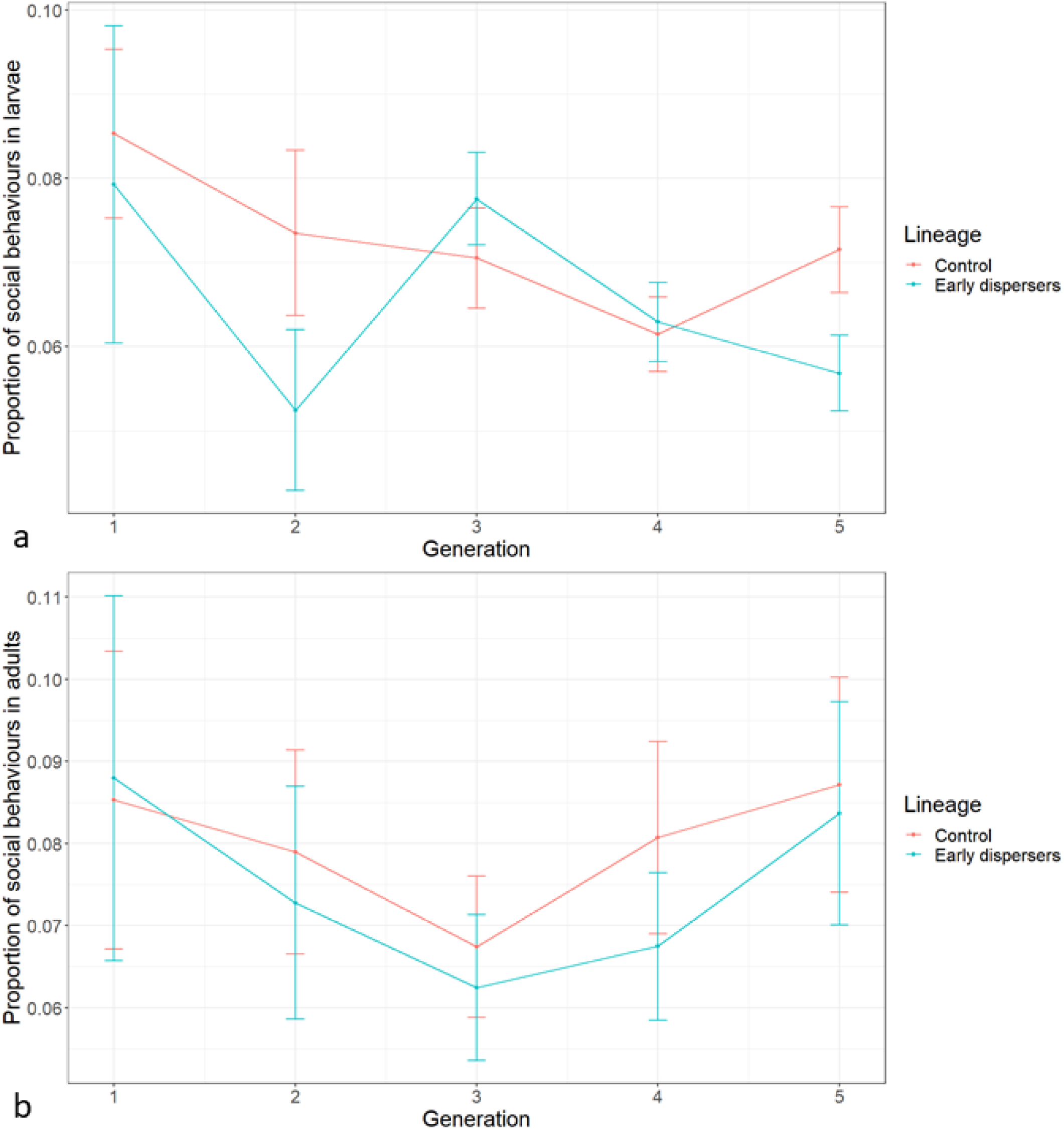
Social behaviors of the two lineages, from F1 to F5 in a) larvae and b) adults. The proportion of social behaviors is expressed as the number of behaviors directed to nest hygiene and allogrooming, out of all behaviors recorded. The comparisons were calculated with Wilcoxon tests including Holm correction for multiple testing. The vertical bars show the standard errors.

Similarly, in both the control and the treatment lineage, the proportion of social behaviors of adults was equal over all consecutive generations (linear models, all p > 0.05). There was no difference between the two lineages in any of the generations (linear models, all p > 0.05) (Fig. 5B).

In the control lineage, the proportion of active behaviors of larvae was higher in F1 than in F5 (control F1 vs. F5, linear model, z = 3.65, p < 0.001) and in F2 (control F1 vs. F2, linear model, z = 7.17, p < 0.001), but lower in F2 than in F3 (control F2 vs. F3, linear model, z = –4.7, p < 0.001). No other two successive generations were found to be significantly different (linear models, all p > 0.05). In the treatment lineage, the proportion of active behaviors of larvae was higher in F1 than in F5 (treatment F1 vs. F5, linear model, z = 4.13, p < 0.001) and in F2 (treatment F1 vs. F2, linear model, z = 3.517, p = 0.02), lower in F3 than in F4 (treatment F3 vs. F4, linear model, z = –4.74, p < 0.001), and higher in F4 than in F5 (treatment F4 vs. F5, linear model, z = 4.01, p = 0.01). There was no difference between the two lineages in any of the generations (linear models, all p > 0.05) (Fig. 6A).

**Figure 6:**
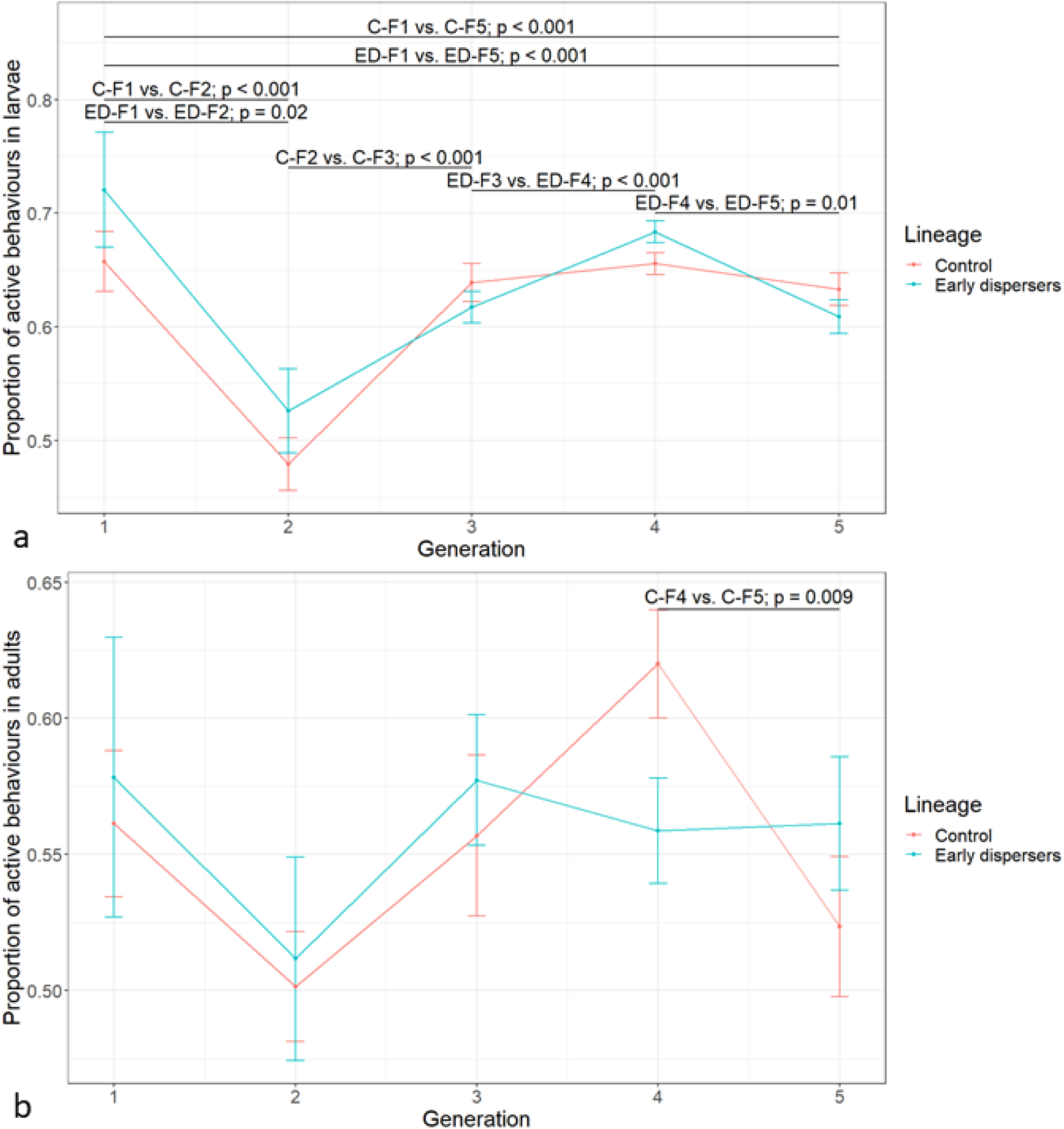
Active behaviors of the two lineages, from F1 to F5 in a) larvae and b) adults. The proportion of active behaviors is expressed as the number of behaviors recorded, excluding the resting behavior. The comparisons were calculated with Wilcoxon tests including Holm correction for multiple testing. The vertical bars show the standard errors.

In the control lineage, the proportion of active behaviors of adults was higher in F4 than in F5 (control F4 vs. F5, linear model, z = 3.64, p = 0.009), and equal over all consecutive generations (linear models, all p > 0.05). In the treatment lineage, the proportion of active behaviors of adults was equal over all consecutive generations (linear models, all p > 0.05). There was no difference between the two lineages in any of the generations (linear models, all p > 0.05) (Fig. 6B).

### Fungal community

The sequencing run of the 65 samples yielded 1203’328 raw reads, with a minimum of 145 reads per sample and a maximum of 34’470. The average number of reads per sample was 18’802. Following alignment and bioinformatics processing, the resulting dataset comprised 59 samples from eight treatment groups, with a total of 1128’177 raw reads. The number of reads per sample ranged from 9’838 to 28’168 (mean = 19’121.64). A total of 70 ASV were identified at the genus level. The water controls yielded a relatively low number of fungal reads, with a mean lower than 218 reads per sample. In contrast, the artificial medium controls yielded a higher number of reads, reaching a level comparable to that observed in the biological samples. The ‘decontam’ package was employed to enhance the quality of the dataset and identified a Sordariomycetes and a Lipomyces ASV as contaminants. Accumulation curves of the final dataset, excluding contaminants and control samples, demonstrated that samples approached saturation after approximately 17 000 reads. Visualization of the relative abundance of the genus revealed that groups appeared homogeneous (Fig. 7).

**Figure 7:**
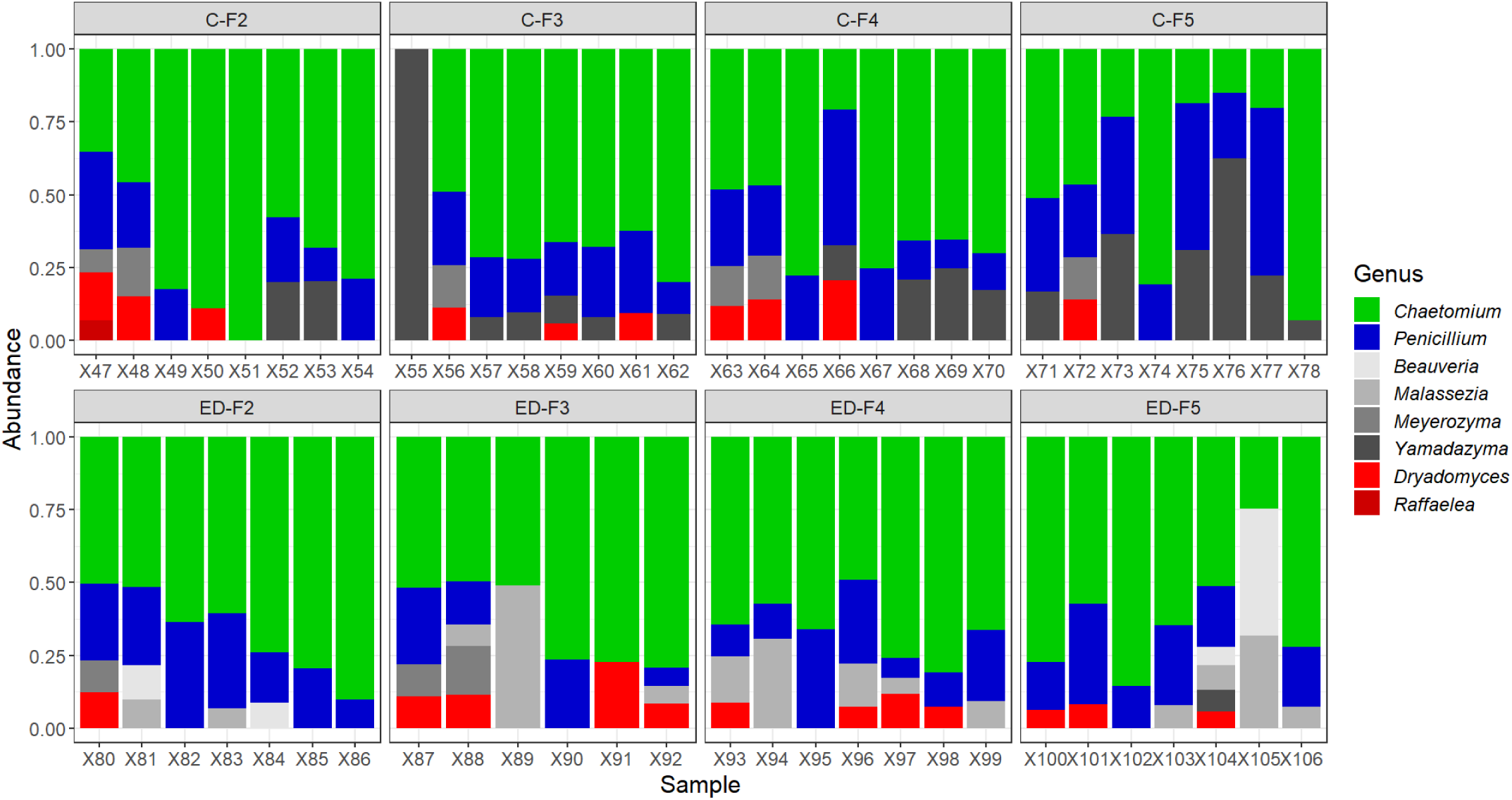
Relative abundances of fungal genera in the two lineages, from F2 to F5. The nutritional symbionts *Dryadomyces* and *Raffaelea* are represented in shades of red, while *Chaetomium* and *Penicillium* are highlighted due to their high abundance. Other fungal genera that are less biologically important are represented in shades of grey.

The Shannon diversity index and observed richness were found to be equal across all generations for each lineage, as well as between lineages for each generation (Wilcoxon test, p > 0.05). (Fig. 8A).

**Figure 8:**
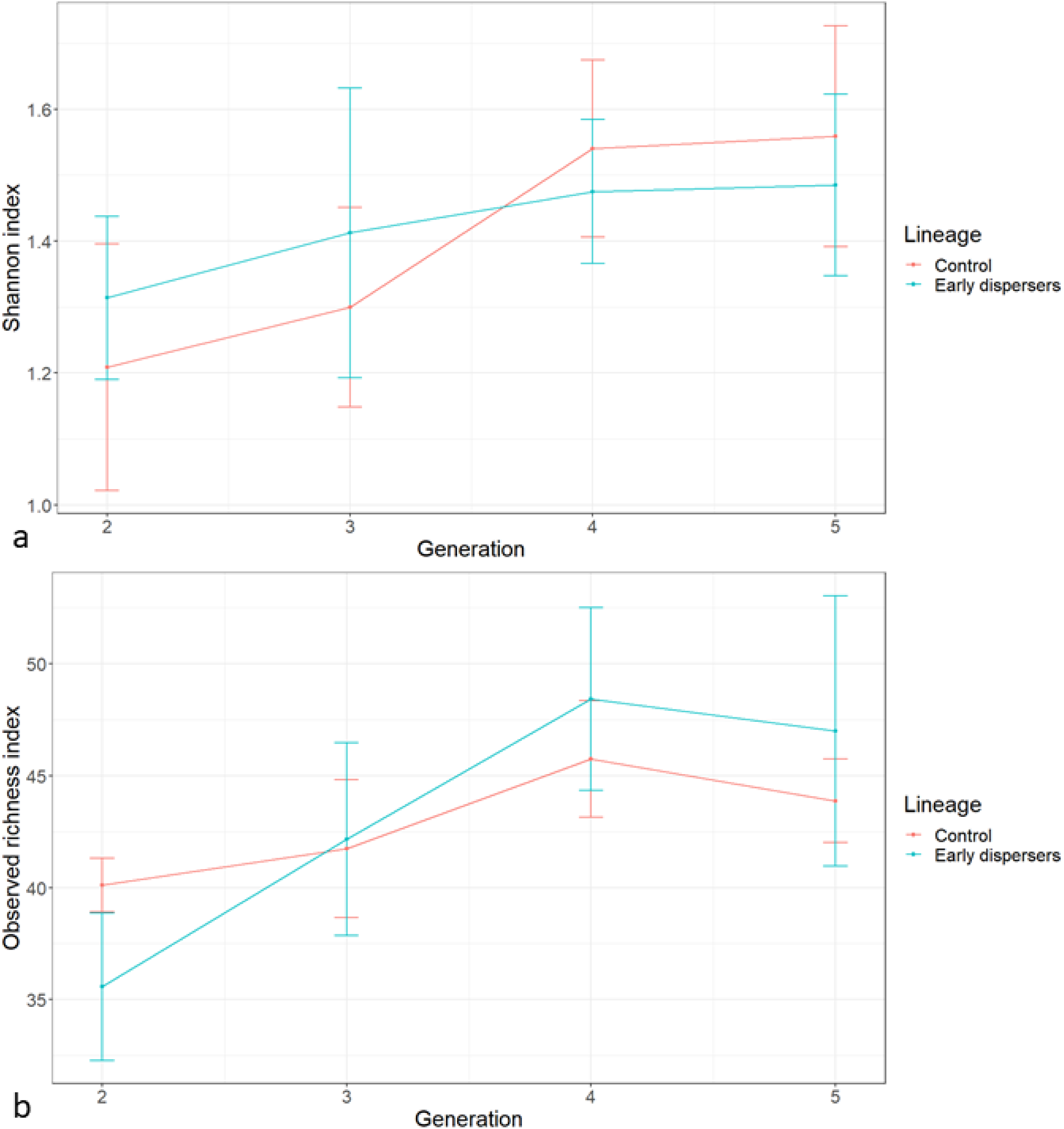
Alpha diversity of the fungal communities of the two lineages, from F2 to F5. a) Shannon index, b) Observed richness. The vertical bars show the standard errors.

In the control lineage, there was a difference in the beta diversity between F2 and F5 (control F2 vs. F5, PERMANOVA, F = 3.36, p = 0.046), but none of the other generations were found to be significantly different (PERMANOVA, all p > 0.05). In the treatment lineage, there was no difference in beta diversity over generations (PERMANOVA, all p > 0.05). A comparison of the two lineages revealed that beta diversities differed in F3, F4 and F5 (control F3 vs. treatment F3, PERMANOVA, F = 1.94, p = 0.044; control F4 vs. treatment F4, PERMANOVA, F = 3.12, p = 0.007; control F5 vs. treatment F5, PERMANOVA, F = 3.93, p = 0.02) (Fig. 7). NMDS visualization of the relative abundance of fungal genera is shown on Fig. 9.

**Figure 9:**
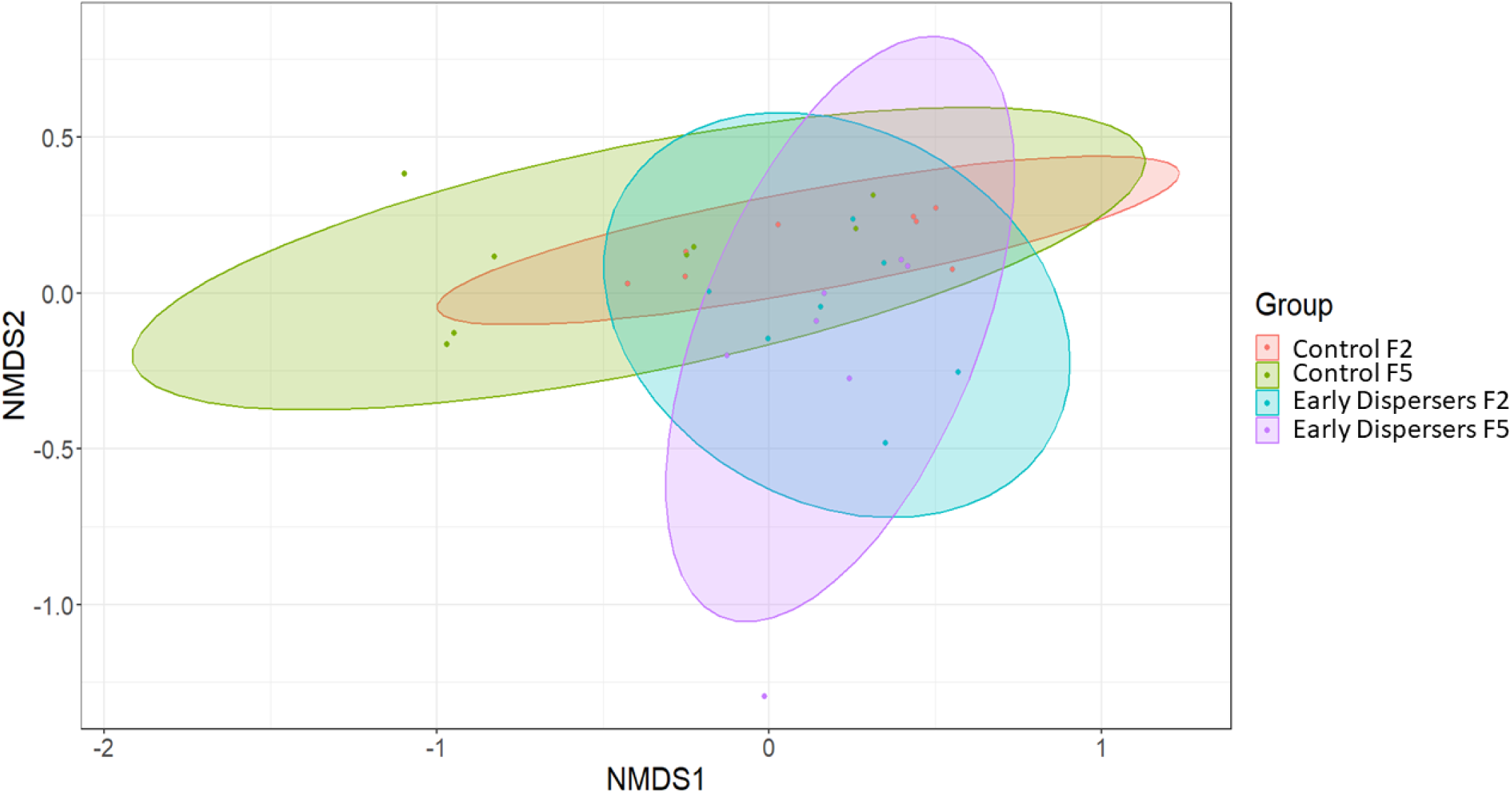
NMDS of the fungal communities of the fungal communities of the two lineages in generations F2 and F5.

## Discussion

The selection experiment was based on pilot results that indicated variable timing of dispersal and correlated effects on the success rate, productivity and social behavior of *X. saxesenii* (Biedermann, 2012). It was hypothesised that this variability could be the result of either inter-individual plasticity or to heritability over generations. The objective of the experiment was to isolate the heritability component by utilising inbred laboratory lineages. The results of the study indicate that the observed variability in the philopatric period is not heritable and can be attributed to inter-individual plasticity alone. Following five generations of selection, no significant responses were identified in the timing of dispersal or the correlated variables, such as social behavior, productivity and the longevity of nests between the control and the treatment lineage. The proportion of social behaviors exhibited by larvae and adults was found to be similar across all groups, indicating that this aspect of *X. saxesenii* behavior is highly conserved. The inbreeding habits of *X. saxesenii* may result in a lack of the necessary variability for an evolutionary response to occur. Prior research has demonstrated that inbreeding and genetic bottlenecks result in a reduction of adaptive potential (Auld & Relyea, 2010; Dierks et al., 2012; Swindell & Bouzat, 2005) and a general decline in phenotypic variance (Fowler & Whitlock, 1999; Reed et al., 2003).

Interestingly, the majority of observed changes in the control lineage occurred between the first and second generations. These findings align with those of other studies that have demonstrated the significant impact of strong selection pressures on the initial generations (Irwin & Carter, 2014; Tejeda et al., 2016). This suggests that the genetic variability present at the onset of the selection experiment, if present, was depleted after the first generation of selection.

At the end of the experiment, the beetles dispersed earlier and exhibited a higher success rate than in the first generation. This indicates that the beetles had become adapted to laboratory conditions. These conditions are favorable to *X. saxesenii*, as the breeding system is maintained at a constant temperature and the artificial medium is rich in nutrients and devoid of competitor microbes. The benign conditions of the laboratory breeding reduce the pressure for cooperation and group living, which consequently favors earlier dispersal. This is consistent with the predictions of theoretical models that have demonstrated a correlation between harsh environments and increased levels of cooperation (Emlen, 1982).

It is worthy of note that a change in the fungal community between beetle lineages was observed over time, with significant differences in beta diversity emerging from the third generation onwards, and significantly increasing in the fourth and fifth generations. This divergence may be attributed to either the selection regime, whereby females dispersing at different times will transmit different fungal communities, or to random drift, whereby the community transmitted by a female in its mycangia does not necessarily represent the complete community developing in the nest. The fungal community associated with *X. saxesenii* is complex, and microbial management is a critical factor in the evolution of cooperative behaviour (Biedermann & Rohlfs, 2017; Nuotclà et al., 2019). The mutualist fungi are conserved and transmitted through generations in the mycetangia (Biedermann et al., 2013; Diehl et al., 2022; Francke-Grosmann, 1975), and are significantly associated with the productivity of *X. saxesenii* (Biedermann et al., 2013; Nuotclà et al., 2021). The results demonstrate that the fungal communities can diverge without significantly impacting *X. saxesenii*, suggesting that the observed difference in beta diversity is not attributed to the nutritional fungi and important antagonists but rather to functionally unimportant fungi that exert no fitness difference in beetles.

In several species of ambrosia beetle, adult females delay dispersal for a variable duration, with not all reproductive adults leaving their native nest at the same time (Biedermann et al., 2011; Nuotclà et al., 2021; Peer & Taborsky, 2007). The results confirmed that the reduction in delay prior to the initial dispersal of adult females of *X. saxesenii* nests was associated with a reduction in lifespan and productivity of the nest. This is experimental evidence demonstrating that delayed dispersal is a major mechanism in the evolution of the social system of *X. saxesenii.* Moreover, the findings revealed that the initial generations demonstrated the most significant response to the selection protocol. To gain insight into the evolutionary potential of this species, future research should focus on investigating the genetic variability present within and between populations of *X. saxesenii*. Subsequent studies should then examine the immediate impact of evolutionary pressures on *X. saxesenii* and investigate the relationship between productivity and the philopatric period in greater detail. Given that *X. saxesenii* is not a conventional model organism, there is still much to be discovered about this system, which provides a unique perspective on the evolution of sociality.

## Notes

https://github.com/AntMelet/Artificial-selection-on-X.-saxesenii

### Competing Interest Statement

The authors have declared no competing interest.

